# Tomato Prf requires NLR helpers NRC2 and NRC3 to confer resistance against the bacterial speck pathogen *Pseudomonas syringae* pv. *tomato*

**DOI:** 10.1101/595744

**Authors:** Chih-Hang Wu, Sophien Kamoun

## Abstract

Bacterial speck, caused by the pathogen *Pseudomonas syringae* pv. *tomato*, is one of the most common diseases in tomato production. Together with Pto kinase, the NLR (nucleotide-binding domain leucine-rich repeat containing) protein Prf confers resistance against the bacterial speck pathogen by recognizing AvrPto and AvrPtoB, two Type III effector proteins secreted by *P. syringae* pv. *tomato*. This Prf/Pto pathway is part of a complex NLR network in solanaceous plants that mediates resistance to diverse pathogens through the helper NLR proteins NRCs (NLR required for cell death). We previously showed that, *in Nicotiana benthamiana*, the hypersensitive cell death elicited by expression of AvrPto and Pto, which activate immunity through the endogenous *Prf* ortholog *NbPrf*, requires functionally redundant *NRC2* and *NRC3*. However, whether tomato (*Solanum lycopersicum*) Prf (*SlPrf*) confers resistance to the bacterial speck pathogen through *NRC2* and *NRC3* has not been determined. In this study, we show that *SlPrf* requires *NRC2* and *NRC3* to trigger hypersensitive cell death and disease resistance in both *N. benthamiana* and tomato. We found that the hypersensitive cell death induced by AvrPtoB/Pto/SlPrf in *N. benthamiana* is compromised when *NRC2* and *NRC3* are silenced, indicating that *SlPrf* is an *NRC2/3*-dependent NLR. We validated this finding by showing that silencing *NRC2* and *NRC3* in the bacterial speck resistant tomato ‘Rio Grande 76R’ compromised Prf-mediated resistance. These results indicate that the NRC network extends beyond *N. benthamiana* to solanaceous crops.

## INTRODUCTION

Plants rely on cell surface receptor and intracellular nucleotide-binding domain leucine-rich repeat (NLR) proteins to mediate pathogen recognition and disease resistance (Dangl et al., 2013; Win et al., 2012). Most plant disease resistance genes encode intracellular NLR receptors that sense pathogen effector proteins directly or indirectly through effector-targeted host proteins (Dangl et al., 2013; Win et al., 2012). An emerging concept in NLR biology is that some “sensor” NLR proteins that detect pathogen molecules require “helper” NLRs, forming complex NLR networks to mediate immune signaling (Bonardi et al., 2011; Wu et al., 2017; Wu et al., 2018). One of such examples is the NLR network mediated by the NRC (NLR-required for cell death) family, in which several solanaceous senor NLR proteins that detect molecules from various pathogens signal through partially redundant NRCs. This NLR network includes several well-known sensor NLRs, such as Rx, Bs2, Sw5b, Mi-1.2 and Rpi-blb2, which confers resistance to virus, bacteria, nematodes and oomycete (Wu et al., 2017). Additionally, the NRCs (NRC2, NRC3 and NRC4) were shown to display functional redundancy and specificity to their matching sensor NLRs (Wu et al., 2017).

Bacterial speck, caused by the hemibiotrophic pathogen *Pseudomonas syringae* pv. *tomato*, is one of the most common bacterial diseases of tomato production (Jones, 1991). It has long been used as a model system for understanding the molecular basis of bacterial pathogenesis and plant disease resistance (Oh and Martin, 2011; Pedley and Martin, 2003). Thus far, the only NLR protein that confers resistance to bacterial speck in tomato is Prf, which forms immune complexes with the intracellular kinase protein Pto. This Pto/Prf complex binds two type III secretion effector proteins from *P. syringae*, AvrPto and AvrPtoB, leading to hypersensitive cell death and disease resistance (Oh and Martin, 2011; Pedley and Martin, 2003). Interestingly, when transiently expressed in *Nicotiana benthamiana*, AvrPto triggers Pto-dependent cell death by signalling through an endogenous Prf ortholog (NbPrf), whereas AvrPtoB triggers Pto-dependent cell death only in the presence of the tomato Prf ortholog (S/Prf) (Balmuth and Rathjen, 2007; Lu et al., 2003; Mucyn et al., 2006).

The immunity pathway induced by Pto/Prf complex is part of a recently described NLR network connected by NRC nodes that are genetically downstream of the sensor NLRs (Wu et al., 2017). We previously showed that, when transiently expressed in *N. benthamiana*, AvrPto and Pto activate *NRC2a* and *NRC2b* (referred to here as *NRC2* for simplicity) as well as *NRC3* dependent cell death, indicating that *NbPrf* requires functionally redundant *NRC2* and *NRC3* to mediate immunity (Wu et al., 2016). However, whether *Prf* displays genetic dependency on *NRC2* and *NRC3* to mediate cell death and disease resistance in tomato has not been determined yet. In this study, we showed that hypersensitive cell death obtained by coexpressing AvrPtoB, Pto and S/Prf in *N. benthamiana* is compromised when *NRC2* and *NRC3* are silenced. Furthermore, the resistance mediated by Pto and Prf in tomato cultivar ‘Rio Grande 76R’ was compromised in *NRC2/3* silenced plants. These results indicate that bacterial speck resistance mediated by Pto/Prf in tomato is *NRC2/3* dependent similar to earlier findings in the *N. benthamiana* system.

## MATERIALS AND METHODS

*N. benthamiana* and tomato plants were grown in a controlled growth chamber with temperature 22-25°C, humidity 45-65% and16/8-h light/dark cycle. Virus-induced gene silencing (VIGS) was performed in *N. benthamiana* and tomato as described by Liu *et al*. (2002). Suspension of *Agrobacterium tumefaciens* strain GV3101 harboring TRV RNA1 (pYL155) and TRV RNA2 (pYL279) (Liu et al., 2002), with corresponding fragments from indicated genes, were mixed in a 2:1 ratio in infiltration buffer (10 mM MES, 10mM MgCl2, and 150 μM acetosyringone, pH5.6) to a final OD600 of 0.3. Two-week-old *N. benthamiana* and tomato plants were infiltrated with *A. tumefaciens* for VIGS inoculations, and then the upper leaves were used two to three weeks later for further agroinfiltrations or disease resistance assays. For silencing of *NRC2/3* homologs in tomato, 5’ coding region of each gene *(S/NRC2*, 1-402bp; *S/NRC3,1-398bp)* were connected together by overlap PCR and cloned into TRV RNA2 vector. Silencing constructs for *NRC2/3* in *N. benthamiana* and *NRC1* in tomato were described previously (Wu et al., 2016)

Transient expression of Prf, Pto, AvrPto and AvrPtoB were performed according to methods described previously (Bos et al., 2006). Briefly, *A. tumefaciens* suspensions were prepared and mixed in the infiltration buffer (10 mM MES, 10 mM MgCl2, and 150 μM acetosyringone, pH5.6). Four to five-week-old *N. benthamiana* plants (i.e. two to three weeks after virus inoculation) were infiltrated with *A. tumefaciens* stains carrying the expression vector of different proteins. The hypersensitive cell death (HR phenotype was imaged at 6 dpi.

Plant total RNA was extracted using RNeasy Mini Kit (Qiagen). First strand cDNA synthesis was performed using SuperScript III Reverse Transcriptase (Thermo Scientific). Semi-quantitative reverse transcription polymerase chain reaction (RT-PCR) was performed using DreamTaq (Thermo Scientific) with 25 to 32 amplification cycles followed by electrophoresis with 1.2% agarose gel stained with ethidium bromide. The following primers were used for RT-PCR: NRC1 (CCAGGAAATTGATCCGCTTA/GATTGCTCACCACCGAATTT), NRC2 (ATGTTGCACGAGTTTTGCAG/GGGTATGGTTGGGATGTCAG), and NRC3 (GATGGAAGCTTGGGATCGTA/GCCCATACATTTTTCGGCTA). Primers for internal control *EF1 a* was described previously (Segonzac et al., 2011).

For testing *Pto/Prf*-mediate resistance to *P. syringae* pv. *tomato* DC3000, the resistant and susceptible tomato cultivars ‘Rio Grande 76R’ and ‘Rio Grande 76S’ were used (Salmeron et al., 1994). Cotyledons of two-week-old tomato seedlings were inoculated with VIGS constructs targeting S7NRC1, S7NRC2 and/or S7NRC3. Inoculation of *P. syringae* pv. *tomato* DC3000 was performed 2-3 weeks after VIGS inoculation according to previous description with minor modifications (Balmuth and Rathjen, 2007). Briefly, *P. syringae* pv. *tomato* DC3000 culture *was* adjusted to OD_600_ of 0.2 and then diluted 10,000-fold with 10mM MgCl2 with 0.02% Silwet L-77. The bacteria suspension was then vacuum-infiltrated into leaves of tomato plants described above. Leaves discs from two plants were sampled using 0.33cm^2^ cork borer at each time points and then homogenized in 10 mM MgCl_2_ for serial dilution and plating. Experiments were repeated three times with similar results.

## RESULTS AND DISCUSSION

We previously showed that AvrPto/Pto induces hypersensitive cell death responses that require the helper NLRs *NRC2* and *NRC3* in *N. benthamiana*. This hypersensitive cell death involves the endogenous sensor NLR protein NbPrf (Lu et al., 2003), leading us to hypothesize that its ortholog in tomato, S7Prf, is also *NRC2/3* dependent. To test this hypothesis, we transiently expressed AvrPto/Pto and AvrPtoB/Pto together with or without S7Prf in *N. benthamiana* leaves. AvrPtoB induced cell death only when S7Prf was co-expressed with Pto consistent with earlier findings (Balmuth and Rathjen, 2007; Mucyn et al., 2006). This cell death response was compromised in the *NRC2/3*-silenced leaves, indicating that S7Prf requires *NRC2* or *NRC3* to induce cell death in *N. benthamiana* (Fig. 1.). Consistent with previous reports (Balmuth and Rathjen, 2007; Mucyn et al., 2006), AvrPto induced cell death regardless of whether or not S7Prf was co-expressed with Pto. This cell death response was also compromised in *NRC2/3*-silenced leaves (Fig. 1.).

**Fig. 1.**
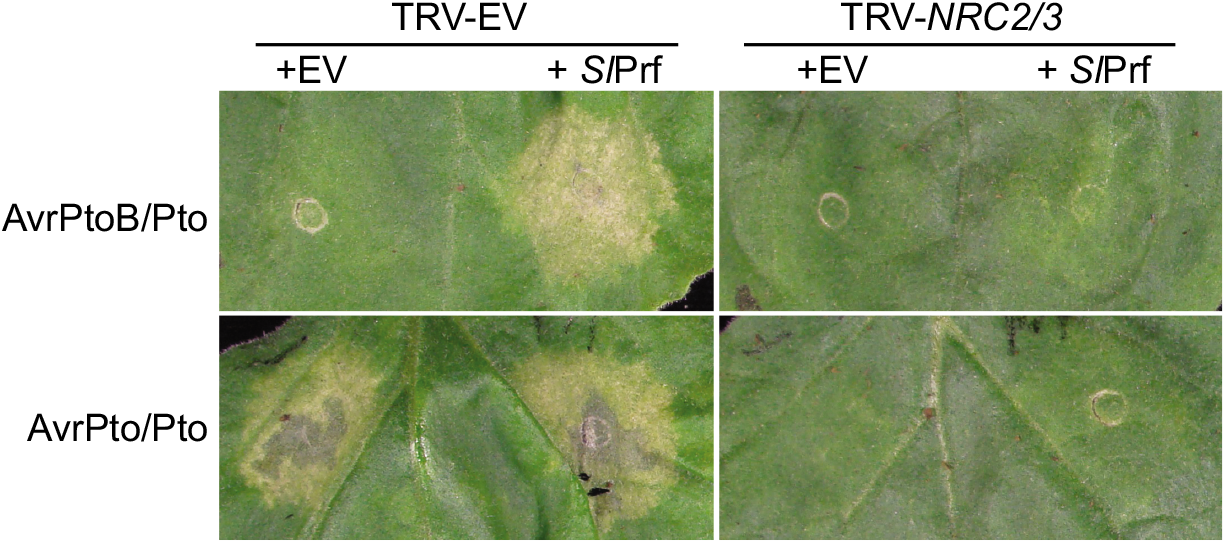
*NRC2* and *NRC3* are required for Prf-mediated cell death in *N. benthamiana*. Pto were co-expressed with AvrPtoB or AvrPto with or without *Sl*Prf in control or *NRC2/3*-silenced *N. benthamiana* leaves. Pictures were taken 5 days after agroinfiltration.

Previously, we used *N. benthamiana* as an experimental system to determine the NRC network architecture (Wu et al., 2017; Wu et al., 2016). We sought out to test whether *NRC2* and *NRC3* are required for *Pto/Prf*-mediated resistance against the bacterial speck pathogen in tomato. We performed disease resistance assays with *P. syringae* pv. *tomato* DC3000 on the tomato bacterial speck resistant cultivar ‘Rio Grande 76R’ and the near-isogenic susceptible cultivar ‘Rio Grande 76R’ We silenced *NRC1, NRC2, NRC3* independently and *NRC2/3* simultaneously using virus-induced gene silencing (Fig. 2A), and then inoculated *P. syringae* pv. *tomato* DC3000 through vacuum infiltration. We observed bacterial speck symptoms in ‘Rio Grande 76S’ plants and ‘Rio Grande 76R’ plants simultaneously silenced for *NRC2* and *NRC3*, but not in ‘Rio Grande 76R’ plants where *NRC1, NRC2*, or *NRC3* were silenced individually (Fig. 2B). Furthermore, bacteria growth assay showed that there was a significantly higher population of bacteria in ‘Rio Grande 76R’ that were co-silenced for *NRC2* and *NRC3* (Fig 2C). Silencing of *NRC1, NRC2* or *NRC3* individually did not affect bacterial growth in tomato ‘Rio Grande 76R’ (Fig. 2C). These results indicate that *NRC2* and *NRC3* are redundantly required for *Pto/Prf-*-mediated disease resistance in tomato (Fig. 3).

**Fig. 2.**
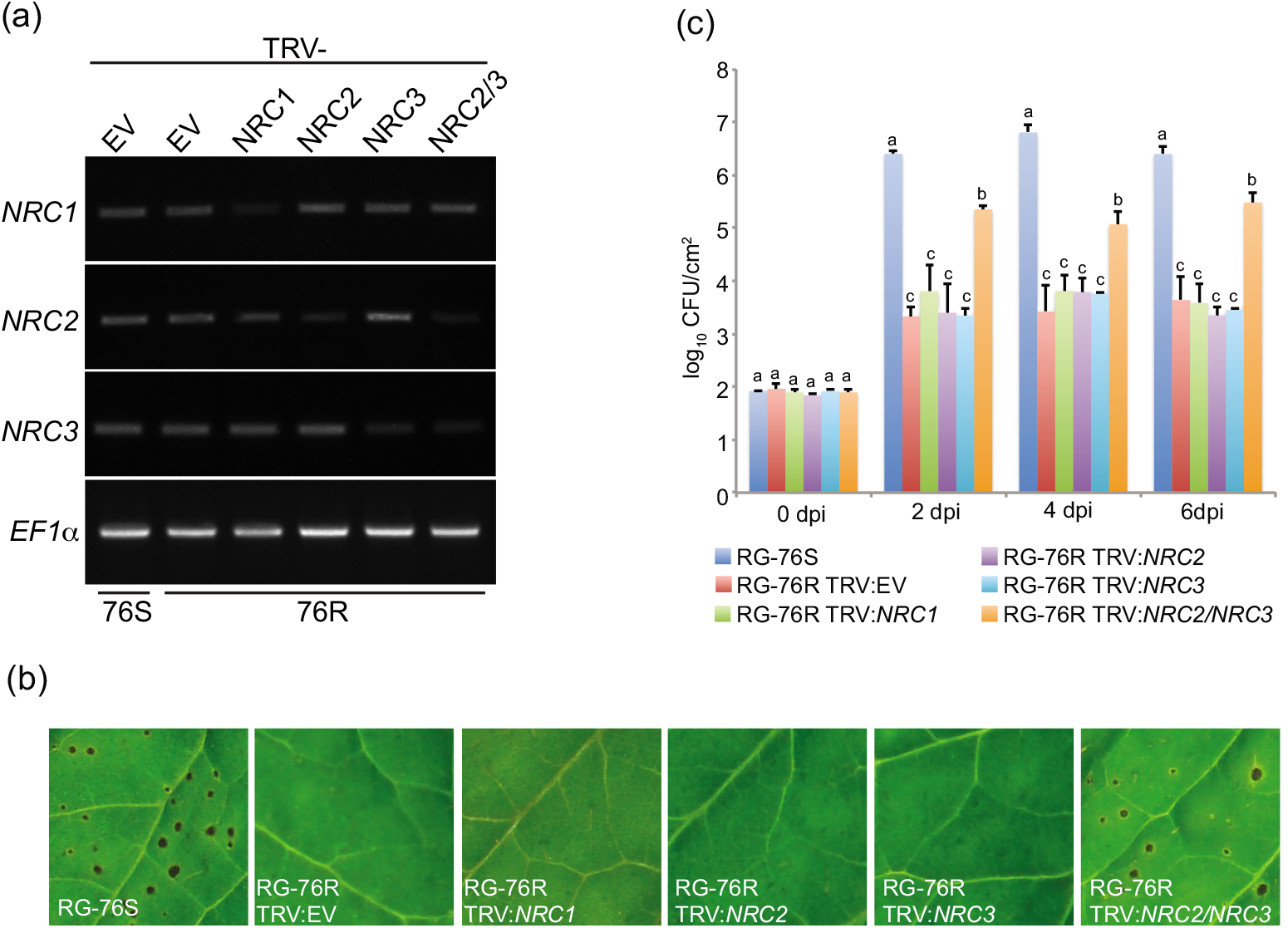
*NRC2* and *NRC3* are required for *Pto/Prf*-mediated resistance in tomato. (a) Semi-quantitative RT-PCR of *NRC*-silenced tomato leaves. Leaves were collected three weeks after virus inoculation. *Elongation factor-1α* (*EFlα*) was used as an internal control. (b) Bacterial speck symptom caused by *P. syringae* on tomato leaves. Pictures were taken 5 days after pathogen inoculation. (c) Bacterial growth assay of *P. syringae* DC3000 in NRC-silenced tomato. Population of *P. syringae* DC3000 were measured at 0, 2, 4, 6 days after inoculation. Error bars indicate the standard deviation of population from one representative biological replicate. Statistical differences among the samples were analyzed with ANOVA and Tukey’s HSD test (p-value < 0.05).

**Fig. 3.**
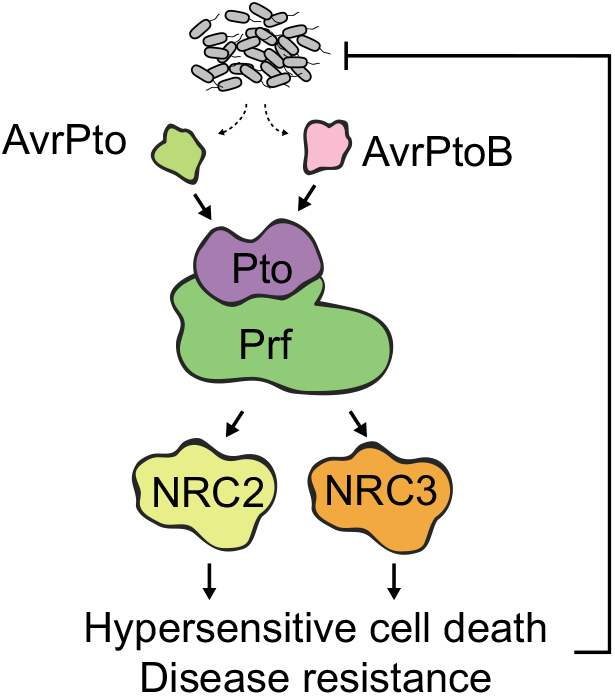
Immunity pathway of Pto/Prf complex in tomato. The kinase protein Pto and the NLR protein Prf form an immune complex that recognizes AvrPto and AvrPtoB secreted from *P. syringae* pv. *tomato*. This complex requires the helper NLR proteins NRC2 and NRC3 to mediated hypersensitive cell death and disease resistance.

However, in the bacterial growth assay, co-silencing of *NRC2* and *NRC3* only partially compromised *Prf*-mediated resistance. This phenomenon could be due to insufficient silencing of the *NRC* genes or the contribution of other tomato *NRCs*, such as *NRC1* that was initially identified as an NLR required for cell death mediated by Prf and other immune receptors (Gabriels et al., 2007). Future experiments are required to determine whether or not *NRC1* or other *NRC* family members are also involved in bacterial speck resistance mediated by *Prf* in tomato. We are currently generating CRISPR/Cas9 mutants of various *NRC* genes that would help address this question.

## CONCLUSION AND PERSPECTIVES

We showed that *NRC2* and *NRC3* are redundantly required for the Pto/Prf immune complex to confer resistance against the bacterial speck pathogen in tomato. These results validate our previous findings with the *N. benthamiana* experimental system that the NLR *Prf* requires the helper NLRs *NRC2* and *NRC3* to mediate immune responses. Future experiments with CRISPR/Cas9 mutants of various *NRC* genes should help us to further address the genetic architecture of NLR networks in tomato and other solanaceous crops.

## ACKNOWLEDGEMENTS

This project was funded by the Gatsby Charitable Foundation, Biotechnology and Biological Sciences Research Council, and European Research Council. We thank Lida Derevnina for carefully reading the manuscript.

